# Molecular mechanism of target site selection and remodeling by type V CRISPR-associated transposons

**DOI:** 10.1101/2021.07.06.451292

**Authors:** Irma Querques, Michael Schmitz, Seraina Oberli, Christelle Chanez, Martin Jinek

## Abstract

Although the canonical function of CRISPR-Cas systems is to provide adaptive immunity against mobile genetic elements^1^, type I-F, I-B and V-K systems have been adopted by Tn7-like transposons to direct RNA-guided transposon insertion^2–7^. Type V-K CRISPR-associated transposons rely on the activities of the pseudonuclease Cas12k, the transposase TnsB, the AAA+ ATPase TnsC and the zinc-finger protein TniQ^7^. However, the molecular and structural details of RNA-directed DNA transposition have remained elusive. Here we report cryo-electron microscopic structures of a Cas12k-guide RNA-target DNA complex and a DNA-bound, polymeric TnsC filament. The Cas12k complex structure reveals an intricate guide RNA architecture and critical interactions mediating RNA-guided target DNA recognition. The assembly of the TnsC helical filament is ATP-dependent and accompanied by structural remodeling of the bound DNA duplex. *In vivo* transposition assays corroborate key features of the structures, and biochemical experiments further show that TniQ restricts TnsC polymerization, while the TnsB transposase interacts directly with TnsC filaments to trigger their disassembly upon ATP hydrolysis. Together, these results suggest a mechanistic model whereby RNA-directed target selection by Cas12k primes TnsC polymerization and DNA remodeling, generating a recruitment platform for TnsB to catalyze site-specific transposon insertion. The present work advances our mechanistic understanding of the cross-talk between CRISPR effectors and the transposition machinery and will inform design efforts to harness CRISPR-associated transposons as programmable site-specific gene insertion tools for genome engineering applications.

## INTRODUCTION

Transposons are mobile genetic elements that pervade all domains of life, thereby shaping the structure, function and evolution of their host genomes^8,9^. Prokaryotes have evolved genome defense mechanisms to restrict parasitic transposition, including adaptive immunity provided by CRISPR-Cas (Clustered Regularly Interspaced Short Palindromic Repeats-CRISPR associated) systems that rely on Cas proteins and CRISPR RNA (crRNA) guides to target invasive genetic elements for nucleolytic degradation ^1,10^. In contrast to the defensive role of canonical CRISPR-Cas systems, several nuclease-deficient type I-F, I-B and V-K systems have been instead co-opted by a distinct group of Tn7-like transposons to direct RNA-guided transposon DNA insertion into specific target sites^2–7^. In these systems, DNA targeting relies on transposon-encoded Cas-RNPs, involving a Cas3-less multisubunit effector complex (termed Cascade) in type I systems^3^ or a single catalytically inactive Cas12k protein in type V-K systems^7^. Cas12k-mediated DNA targeting requires a crRNA guide and a trans-activating CRISPR RNA (tracrRNA), and is coupled to the concerted activities of three Tn7-like transposon proteins: the transposase TnsB, the AAA+ ATPase TnsC and the zinc-finger protein TniQ to direct unidirectional insertion of the transposon DNA at a fixed distance (60-66 bp) downstream of the target site. In the prototypical *Escherichia coli* Tn7 transposon^11^, the DDE-type TnsB transposase catalyses the 5’-DNA strand breakage and joining reactions required for transposon DNA excision from a donor locus and integration into target DNA^12,13^. The transposition regulator TnsC helps recruit TnsB to the target site and stimulates its transposase activity^14–16^. The Tn7 core transposition machinery lacks an intrinsic mechanism for target site recognition and instead relies on DNA-binding cofactors TnsD and TnsE to direct transposition to specific attachment site sequences^17^ and DNA structures^18,19^, respectively. In CRISPR-associated transposons, TnsD and TnsE are replaced by the crRNA-guided effector (Cascade or Cas12k) and a putative DNA binding protein TniQ. Although the DNA targeting mechanism of type I-F3 CRISPR-transposon systems has been elucidated by structural studies of Cascade-TniQ complexes^20–23^, the structural basis of RNA-guided DNA targeting in type V-K systems remains unknown. Harnessing crRNA-guided transposition would enable programmable DNA insertion without the need for double-strand break-induced homologous recombination. In this context, the current lack of mechanistic information on Cas12k-directed transposition hampers engineering these systems for genome editing applications. To understand the molecular mechanisms underpinning target DNA recognition and transpososome assembly in type V-K CRISPR-associated transposons, we carried out structural studies of the *Scytonema hofmanni* ShCAST system^7^. Here we describe cryo-electron microscopy structures of a ternary Cas12k-guide RNA-target DNA complex and DNA-bound TnsC.

## RESULTS

### Target DNA recognition mechanism of the Cas12k pseudonuclease

To gain insights into how Cas12k recognizes its DNA target, we analyzed using single-particle cryo-electron microscopy (cryo-EM) a ternary complex comprising ShCas12k, single-guide RNA (sgRNA, a 254-nucleotides crRNA-tracrRNA fusion containing a 24-nucleotide guide segment) and target dsDNA (50 base pairs), obtaining a reconstruction at an overall resolution of 3.0 Å (**Fig. 1, Extended Data Fig. 1,2, Supplementary Table 1**). Three-dimensional classification and refinement of recorded particles revealed that the ternary ShCas12k complex exhibits considerable conformational heterogeneity (**Extended Data Fig. 2**). While the majority of sgRNA nucleotides could be modeled, only parts of the R-loop structure formed by the sgRNA and target dsDNA could be resolved in the reconstructed map, corresponding to a 9-base pair duplex of the target (TS) and non-target DNA strands (NTS) containing the protospacer adjacent motif (PAM), and a downstream 9-base pair heteroduplex formed by the sgRNA and PAM-proximal region of the TS DNA. This may be due to the overall conformational heterogeneity of the ternary complex and reflect a situation in which the R-loop is dynamic or only partially formed.

**Fig. 1.**
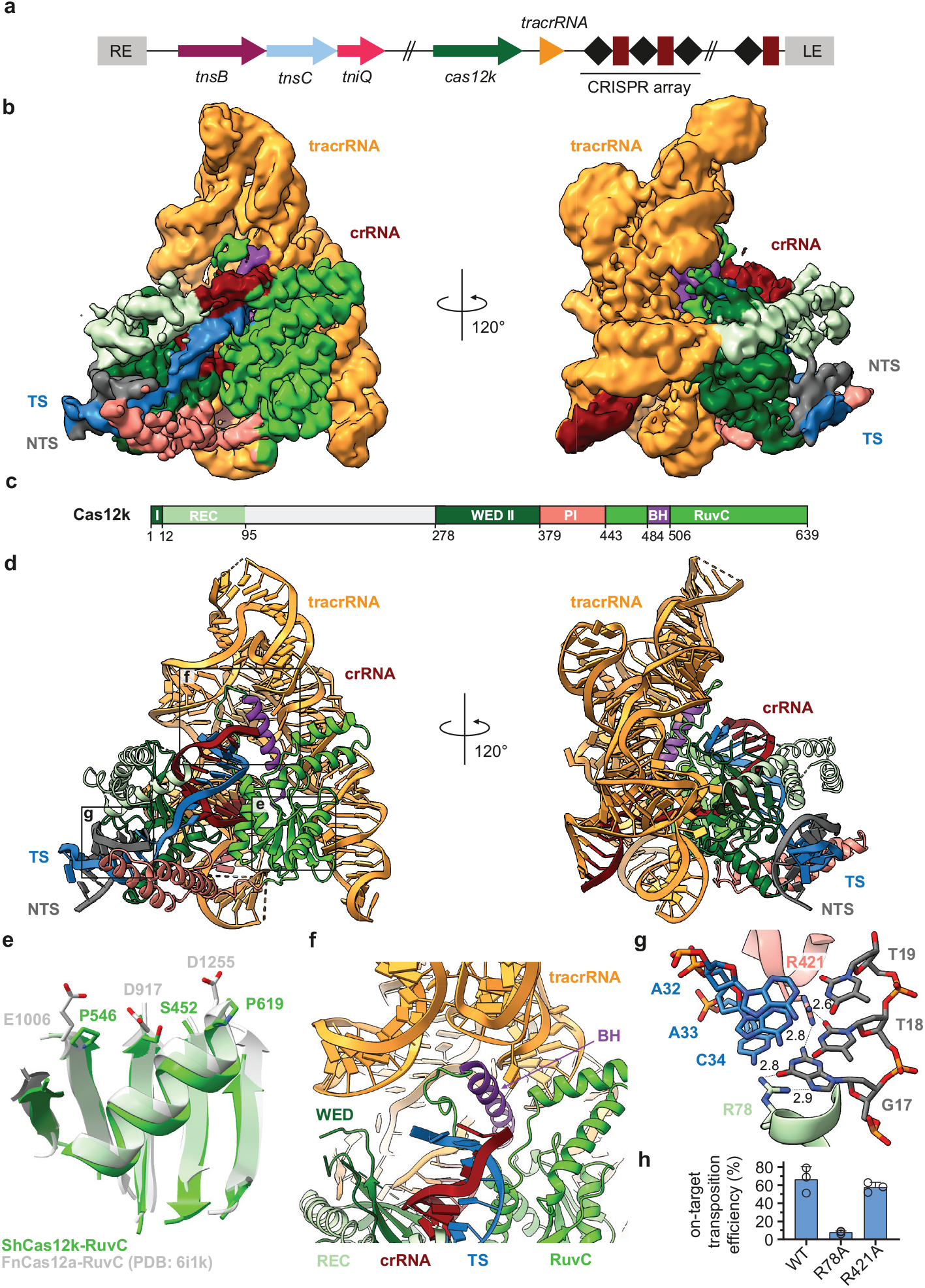
Cryo-EM structure of the ShCAST Cas12-sgRNA-target DNA complex. **a**, Schematic diagram of the type V-K CRISPR-associated transposon system from *Scytonema hofmanni* (UTEX B2349). LE, RE - left and right transposon ends. **b**, Electron density map of the ShCas12k-sgRNA-target DNA complex. TS, target DNA strand; NTS, non-target DNA strand. **c**, Domain organization of ShCas12k. REC, recognition lobe. WED I and II, wedge domain. PI, PAM interacting domain. BH, bridging helix. **d**, Structural model of the ShCas12k-sgRNA-target DNA complex. **e**, Zoomed-in view of the inactivated ShCas12k RuvC domain overlaid with the FnCas12a-RuvC domain (PDB: 6i1k)^53^. **f**, Zoomed-in view of RNA:DNA duplex in proximity of BH. **g**, Details of PAM recognition. Hydrogen bonds are shown as dotted lines. Distances are in Å **h**, Site-specific transposition activity in *E. coli* of ShCAST systems containing mutations in PAM interacting residues, as determined by digital droplet PCR (ddPCR) analysis. Data are represented as mean ± SD (*n*=3).

The general domain architecture of Cas12k is similar to those of other type V CRISPR-associated Cas12 effectors and comprises WED, REC1, REC2, PI (PAM interacting) and RuvC domains (**Fig. 1c,d**). The RuvC domain adopts the canonical fold and superimposes closely with the corresponding domains of other Cas12-family proteins (**Fig. 1e**). However, the RuvC active site is degenerate in Cas12k as it lacks the canonical Asp-Glu-Asp motif involved in Mg^2+^ binding, instead featuring Ser452^ShCas12k^, Pro546^ShCas12k^ and Pro619^ShCas12k^ (**Fig. 1e**), providing a structural confirmation that Cas12k is a pseudonuclease incapable of cleaving target DNA.

In the captured conformational state of the Cas12k ternary complex, the sgRNA-TS DNA heteroduplex is nested in a cleft formed by the REC1 and RuvC domains and terminates after nine base-pairs as it contacts the Cas12k bridge helix (**Fig. 1f**). This suggests that full sgRNA-TS DNA hybridization requires additional conformational rearrangements in Cas12k. Of note, incomplete R-loop formation was previously observed in the structure of the Cascade-TniQ complex from the *Vibrio cholerae* type I-F CRISPR-transposon system^20^, implying that this may be a common feature of CRISPR-associated transposon systems and hinting that full R-loop formation may occur upon subsequent recruitment of the downstream transposon proteins, notably TnsC.

crRNA-guided DNA recognition in the ShCAST system is dependent on the presence of a 5’-GTN-3’ PAM upstream of the target site^7^. In the ternary complex structure, the PAM-containing DNA duplex is contacted by the WED, REC1 and PI domains. The G-C base pair at the −3 PAM position is specifically read out by a bidentate hydrogen bonding interaction of the side chain of Arg78^ShCas12k^ with the major groove edge of the guanine base (**Fig. 1g**). Additionally, Arg421^ShCas12k^ contributes to PAM recognition by contacting the minor groove edges of the guanine and thymine bases at the −3 and −2 PAM positions, respectively, by hydrogen bonding interactions (**Fig. 1g**). To verify the PAM recognition mechanism of Cas12k, we utilized a droplet-digital PCR (ddPCR) assay to quantify the effect of Cas12k mutations on the efficiency of ShCAST-mediated DNA transposition in *Escherichia coli*. Although the R421A mutation resulted in a slight reduction in transposition frequency, the R78A mutation nearly completely abolished transposition *in vivo* (**Fig.1h**), indicating that Arg78^ShCas12k^ is the major structural determinant of PAM recognition in ShCAST.

### An extensive tracRNA scaffold supports Cas12k-dependent transposition

With a length of >250 nucleotides and a molecular weight of 82 kDa, the ShCas12k sgRNA is the largest CRISPR-associated guide RNA scaffold described to date. The RNA has a highly complex pseudoknot topology containing two long coaxial duplex stacks that clamp the Cas12k RuvC domain and a smaller side stack that contains the crRNA-tracrRNA repeat/anti-repeat duplex (**Fig. 2a,b**). The central element of the sgRNA is a triplex region involving nucleotides G226-A229 derived from the crRNA repeat sequence, which contacts the WED domain and connects the repeat/anti-repeat duplex of the sgRNA with the sgRNA-TS DNA duplex (**Fig. 2c**). The central triplex is conserved in the Cas12e (CasX) sgRNA, which is, however, otherwise structurally divergent from the Cas12k sgRNA scaffold (**Extended Data Fig. 3**). Other key structural features of the sgRNA include: a triplex region involving the 5’-terminal segment of the tracrRNA (nucleotides A6-A9) forming A-minor interactions with the pseudoknot duplex stack; a pseudoknot duplex between nucleotides G10-C13 and G143-C146; a “rooftop” stem-loop and a “connector” duplex that bridge the pseudoknot and central stacks; and a triplex junction featuring ribose-zipper interactions between sgRNA nucleotides C96-C97 and A190-A191 (**Fig. 2a,b, Extended Data Fig. 4**). By making extensive interactions with the Cas12k WED and RuvC domains, the tracrRNA plays a major scaffolding role to ensure their correct orientation and present the guide segment of the sgRNA for base-pairing interactions with the target DNA.

**Fig. 2.**
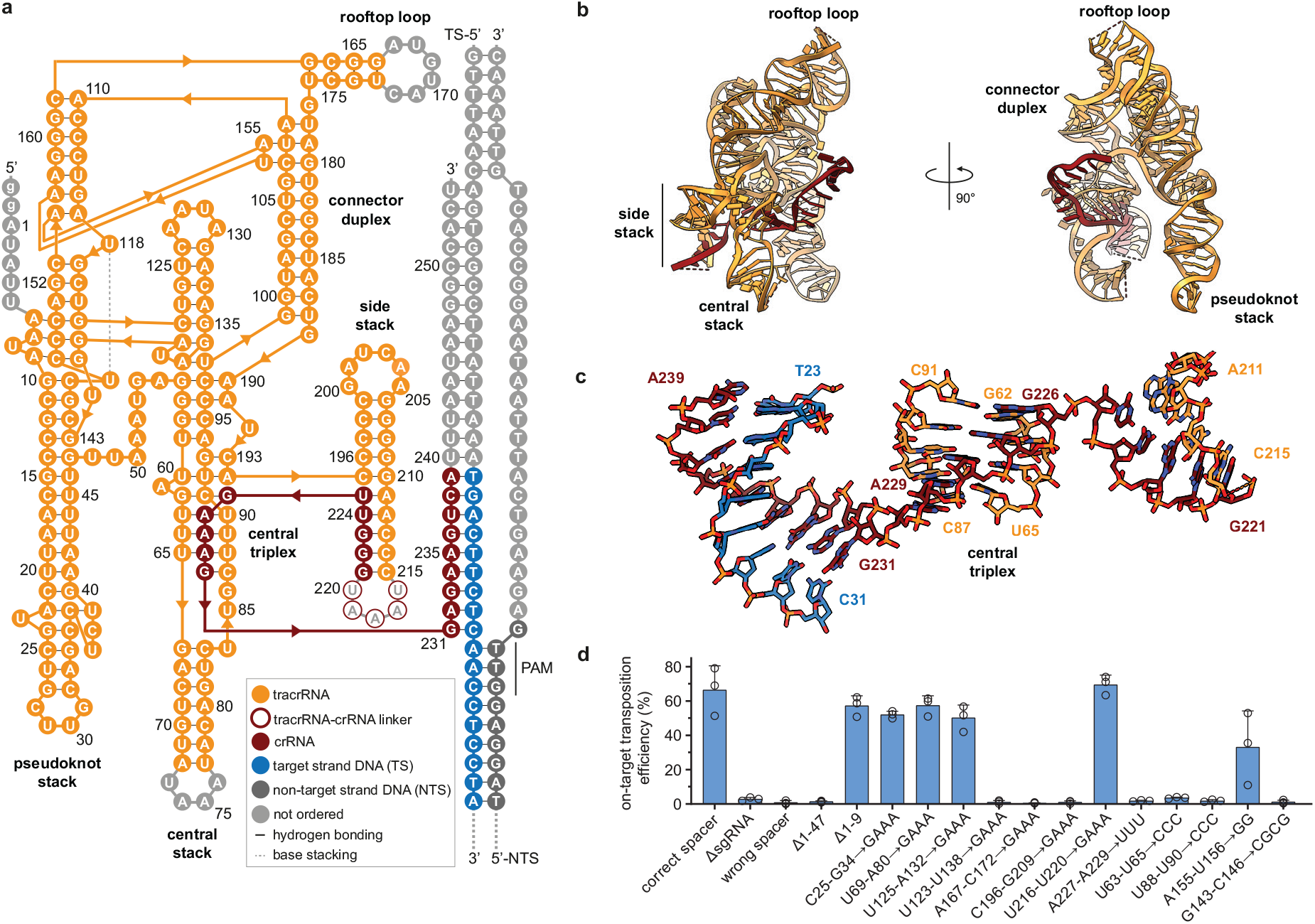
Extensive sgRNA architecture supports Cas12k function in ShCAST. **a**, Schematic diagram of the secondary and tertiary structures of the sgRNA scaffold. **b**, Structural model of the ShCas12k sgRNA. **c**, Zoomed-in view of the sgRNA central triplex. **d**, ddPCR-based transposition activity assay of structure-based sgRNA scaffold mutants of the ShCAST system. Data are represented as mean ± SD (*n*=3).

Using the ddPCR transposition assay, we tested the effect of structure-based sgRNA mutations on transposition efficiency *in vivo* (**Fig. 2d**, **Extended Data Fig. 4f**). Mutations of the central triplex (nucleotides U88-U90), rooftop loop (A167-C172) and and the pseudoknot duplex (G143-C146) truncation of the 5’-terminal region of the sgRNA (nucleotides 1-47) or all resulted in substantial reduction or complete loss of transposition (**Fig. 2d**). In contrast, substitution of the peripheral stem-loops of the sgRNA scaffold (nucleotides C25-G34, U69-A80, U125-A132, U216-U220) with GAAA tetraloops did not perturb transposition efficiency (**Fig. 2d**). These results confirm the functional importance of the key sgRNA structural elements in supporting the transposition activity of ShCAST and suggest ways to minimize the size of the sgRNA scaffold for genome engineering applications.

### The AAA+ ATPase TnsC forms helical filaments

The TnsC component of type V CRISPR-associated transposons is a putative AAA+ ATPase that has been postulated to link Cas12-mediated target site recognition to transpososome recruitment^7,24^. Transposon-associated AAA+ ATPases such as MuB and IstB have previously been shown to oligomerize in the presence of ATP^25,26^. To determine whether this also applies to TnsC, we used negative-stain electron microscopy to directly observe oligomerization of purified ShTnsC. Upon incubation with a linear double-stranded DNA oligonucleotide in the presence of ATP or the non-hydrolyzable analog ATPγS, ShTnsC formed polymeric filaments with an average length of 90 nm. Filament formation did not occur in the absence of dsDNA or ATP (**Fig. 3a, Extended Data Fig. 5**). Together, these results indicate that TnsC polymerization is a DNA-dependent process that requires ATP binding but not hydrolysis, in good agreement with the multimerization properties of other transposon-associated AAA+ ATPases.

**Fig. 3.**
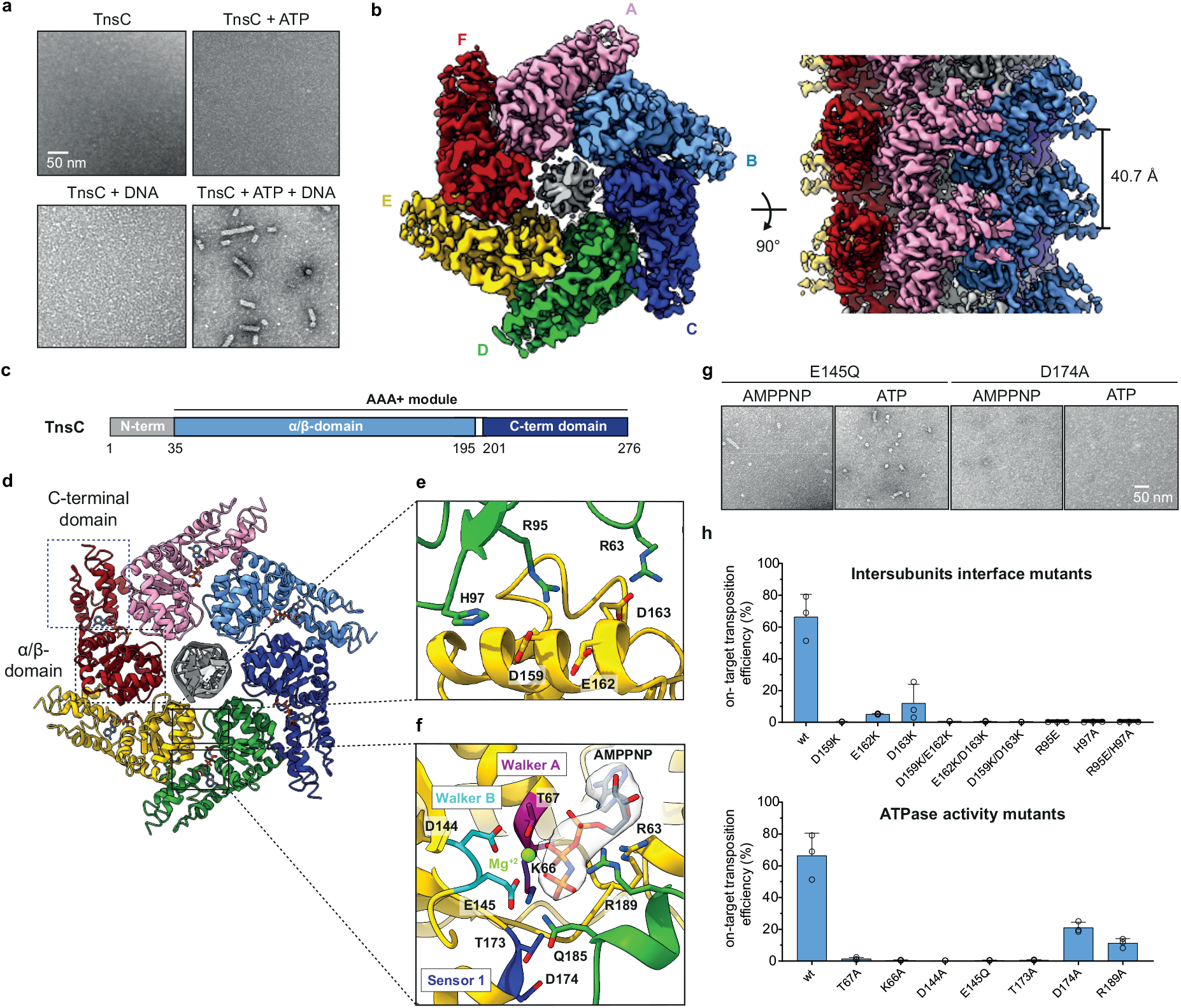
Cryo-EM structure of the ShTnsC•AMPPNP•dsDNA filaments. **a**, Representative negative stain electron micrographs of ShTnsC in the presence or absence of a 92 bp dsDNA and ATP. Scale bars, 50 nm. Magnification, 68,000 x. **b**, Cryo-EM density map of the dsDNA- and AMPPMP-bound ShTnsC filament at 3.6 Å overall resolution (top and side views). Six depicted protein subunits are uniquely colored. dsDNA is shown in grey. **c**, Domain organization of ShTnsC. **d**, Structural model of the ShTnsC filament. **e–f**, Zoomed-in views of the inter-protomer interface and of the ATPase catalytic site. Density corresponding to the AMPPNP nucleotide is shown (contour level of 4.0 σ in Chimera). The AMPPNP molecule and selected protein side chains are shown in stick representation. **g**, Representative negative stain EM micrographs of ShTnsC mutants in the presence of dsDNA and ATP or AMPPNP. Scale bars, 50 nm. Magnification, 98,000 x. **h**, Site-specific transposition activity in *E. coli* of ShCAST systems containing mutations at the inter-protomer interface and in the catalytic pocket of ShTnsC, as determined by digital droplet PCR (ddPCR) analysis. Data are represented as mean ± SD (*n*=3).

We next used cryo-EM to determine the structure of a DNA-bound ShTnsC filament in the presence of the non-hydrolyzable analog AMPPNP and Mg^2+^, at an overall resolution of 3.6 Å, using helical reconstruction (**Supplementary Table 2, Extended Data Fig. 6,7**). The structure reveals that TnsC forms a right-handed helix with a pitch of 40.7 Å and rise of 6.8 Å, perfectly accommodating six ShTnsC copies per turn (**Fig. 3b**). The ShTnsC protomers contain a canonical two-domain AAA+ ATPase module, composed of an α/β domain comprising a five-stranded beta sheet flanked by alpha helices on both sides, and an α-helical C-terminal domain (**Fig. 3c,d, Extended Data Fig. 8**). The N-terminal region of ShTnsC (residues Asp17-Lys31^ShTnsC^) forms and additional alpha helix that co-folds with the C-terminal domain into a single structural unit. The ATPase active site is located in a cleft between the α/β and C-terminal domains and contains the canonical AAA+ structural motifs involved in ATP binding and hydrolysis, including Walker A and Walker B (**Fig. 3f**). The active site is completed by the insertion of the arginine finger Arg189^ShTnsC^ and Gln185^ShTnsC^ from the adjacent ShTnsC protomer, thereby coupling ATP binding to ShTnsC polymerization. The coordination of the AMPPNP ligand and the Mg^2+^ cofactor appears to reflect the pre-hydrolytic state. Helical assembly of ShTnsC protomers is further enabled by additional salt-bridge interactions involving Arg63^ShTnsC^, Arg95^ShTnsC^ and His97^ShTnsC^ from one protomer and Asp159^ShTnsC^, Asp163^ShTnsC^ and Glu162^ShTnsC^ from another (**Fig. 3e**).

To validate our structural findings, we analyzed the polymerization behavior of ShTnsC mutant proteins (**Extended Data Fig. 9a**) by negative-stain EM. The E145Q Walker B mutant protein, predicted to be deficient in ATP hydrolysis but not ATP binding, supported filament formation in the presence of ATP or AMPPNP (**Fig. 3g**). In contrast, ShTnsC containing the D174A mutation in the Sensor 1 motif was incapable of filament formation, confirming that ATP binding but not its hydrolysis is required for ShTnsC assembly. We additionally probed whether ShTnsC polymerization is required for transposition *in vivo* using the ddPCR transposition assay. This experiment revealed that mutations of key residues involved in ATP binding and hydrolysis and in ShTnsC polymerization resulted in a substantial reduction or complete loss of transposition (**Fig. 3h**). Together, these results confirm that both ATP-dependent TnsC oligomerization and ATP hydrolysis by TnsC are critical for transposition activity of type V CRISPR-associated transposon systems.

### TnsC filaments assemble on structurally remodeled DNA

The helical filament generated by ShTnsC completely encloses the bound DNA. Close inspection revealed that the DNA duplex is structurally distorted to match the helical symmetry of the ShTnsC filament. Instead of adopting regular B-form geometry, the DNA duplex has 12 base pairs per helical turn with a pitch of 40.7 Å, a rise of 3.4 Å per base pair along the helical axis, and a narrower diameter (17.6 Å), as compared to a canonical B-form duplex (20 Å) (**Fig. 4a**). The right-handed ShTnsC filament makes continuous interactions with the deoxyribose-phosphate backbone of only one of the two DNA strands (the tracking strand), with each protomer straddling the minor groove of the duplex and contacting two consecutive backbone phosphate groups on the tracking strand via a salt bridge with Lys103^ShTnsC^ and a hydrogen bonding interaction with Thr121^ShTnsC^ (**Fig. 4b**). Although the other DNA strand is not contacted directly by ShTnsC, its backbone phosphate groups are adjacent to Lys99^ShTnsC^ and Lys150^ShTnsC^ (**Fig. 4b**). T121A and K99A ShTnsC mutant proteins generated substantially shorter filaments, while filament formation was not perturbed by the K103A mutation (**Extended Data Fig. 9b, Fig. 4c**). Corroborating these results, the DNA binding activities of T121A, and K99A ShTnsC mutant proteins, as monitored by an electrophoretic mobility shift assay, were also substantially reduced (**Extended Data Fig. 9b, c**). Finally, ShTnsC mutations resulting in defective TnsC polymerization also abolished or significantly reduced DNA transposition *in vivo* (**Fig. 4d**). Together, these results validate the functional importance of TnsC-DNA interactions in the transposition mechanism of ShCAST, implying that DNA-dependent TnsC oligomerization and TnsC-induced DNA remodeling are required for the transposition activity of type V CRISPR-associated transposons.

**Fig. 4.**
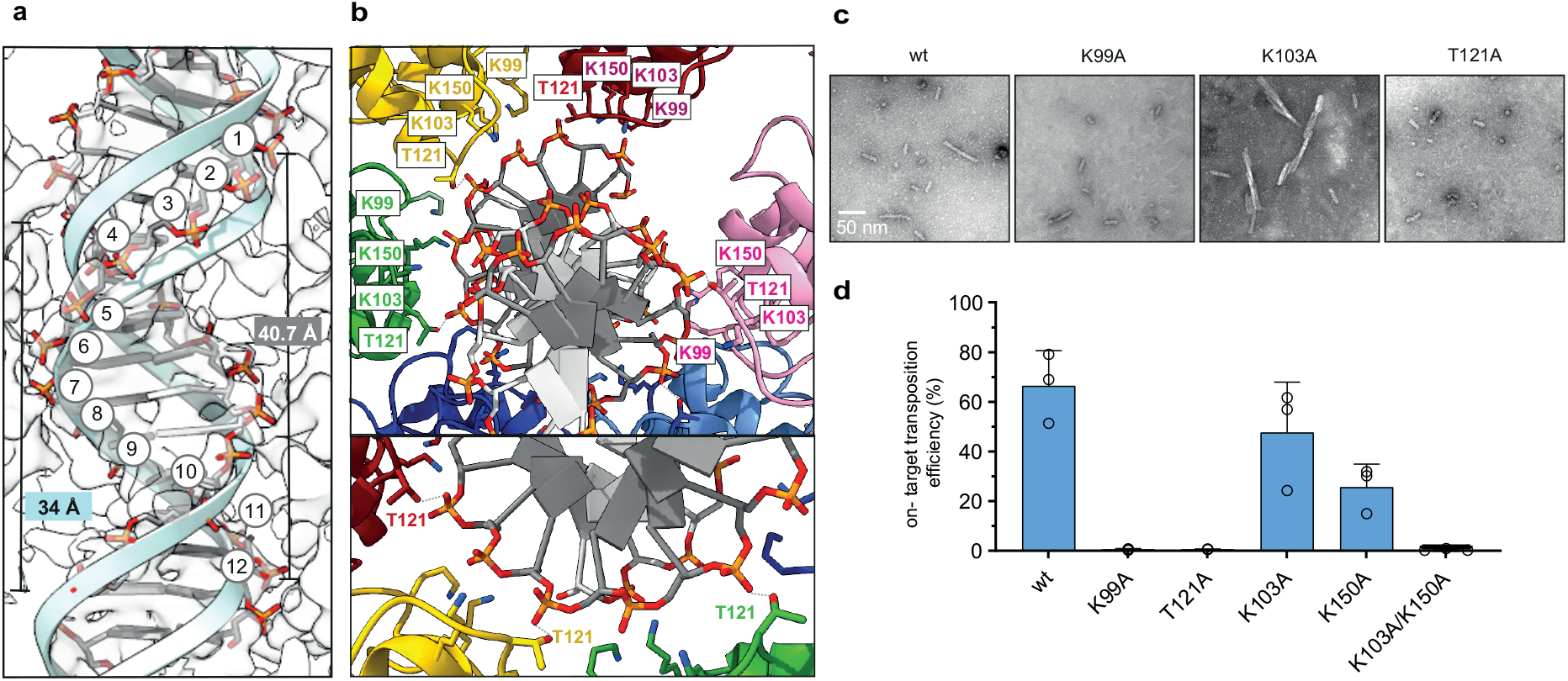
dsDNA binding and remodeling by the ShTnsC filament. **a**, Overview of the bound DNA duplex (backbone in stick representation; tracking strand, dark grey; opposite strand, white) and comparison with an ideal B form dsDNA (light blue cartoon backbone). Backbone phosphates of the tracking strand are numbered. **b**, Close-up views of key DNA-interacting residues (side chains in stick representation). Protein protomers are shown in distinct colors. Hydrogen bonds are indicated with grey dashed lines. **c**, Representative negative stain EM micrographs of ShTnsC in the presence of AMPPNP and dsDNA and with mutations in the DNA binding interface. Scale bars, 50 nm. Magnification, 98,000 x. **d**, Site-specific transposition activity in *E. coli* of ShCAST systems containing mutations in the DNA binding interface of ShTnsC, as determined by digital droplet PCR (ddPCR) analysis. Data are represented as mean ± SD (*n*=3).

### TnsB recruitment by TnsC triggers filament disassembly

In Tn7 and Mu transposons, the transposases TnsB and MuA interact with and stimulate the ATPase activity of their cognate AAA+ ATPases (TnsC^27^ and MuB^28,29^), respectively. To test whether this also occurs in ShCAST, we performed pull-down experiments using affinity-tagged ShTnsB. In the presence of DNA, ShTnsB efficiently co-precipitated ShTnsC when ATP or AMPPNP was also present, but not in the presence of ADP (**Fig. 5a**). A truncated ShTnsB protein comprising the catalytic and C-terminal domains was sufficient to mediate the interaction; however, a TnsB construct lacking the C-terminal domain (residues Leu494-Phe584^ShTnsB^) was unable to co-precipitate ShTnsC. These results indicate that ShTnsB interacts with ShTnsC filaments but not with monomeric ShTnsC, and that the C-terminal region of ShTnsB is required for ShTnsC interaction. We next used negative-stain EM to visualize the effect of ShTnsB on ShTnsC filaments. We observed disassembly of ATP-bound ShTnsC filaments in the presence of ShTnsB (**Fig. 5b**), while AMPPNP-bound filaments remained intact, indicating that effect of ShTnsB was dependent on ATP hydrolysis. Taken together, these results suggest that the C-terminal domain of ShTnsB interacts with and stimulates the ATPase activity of polymerized ShTnsC to trigger filament disassembly, and confirm that transposase-dependent TnsC disassembly is a conserved feature of the ShCAST system.

**Fig. 5.**
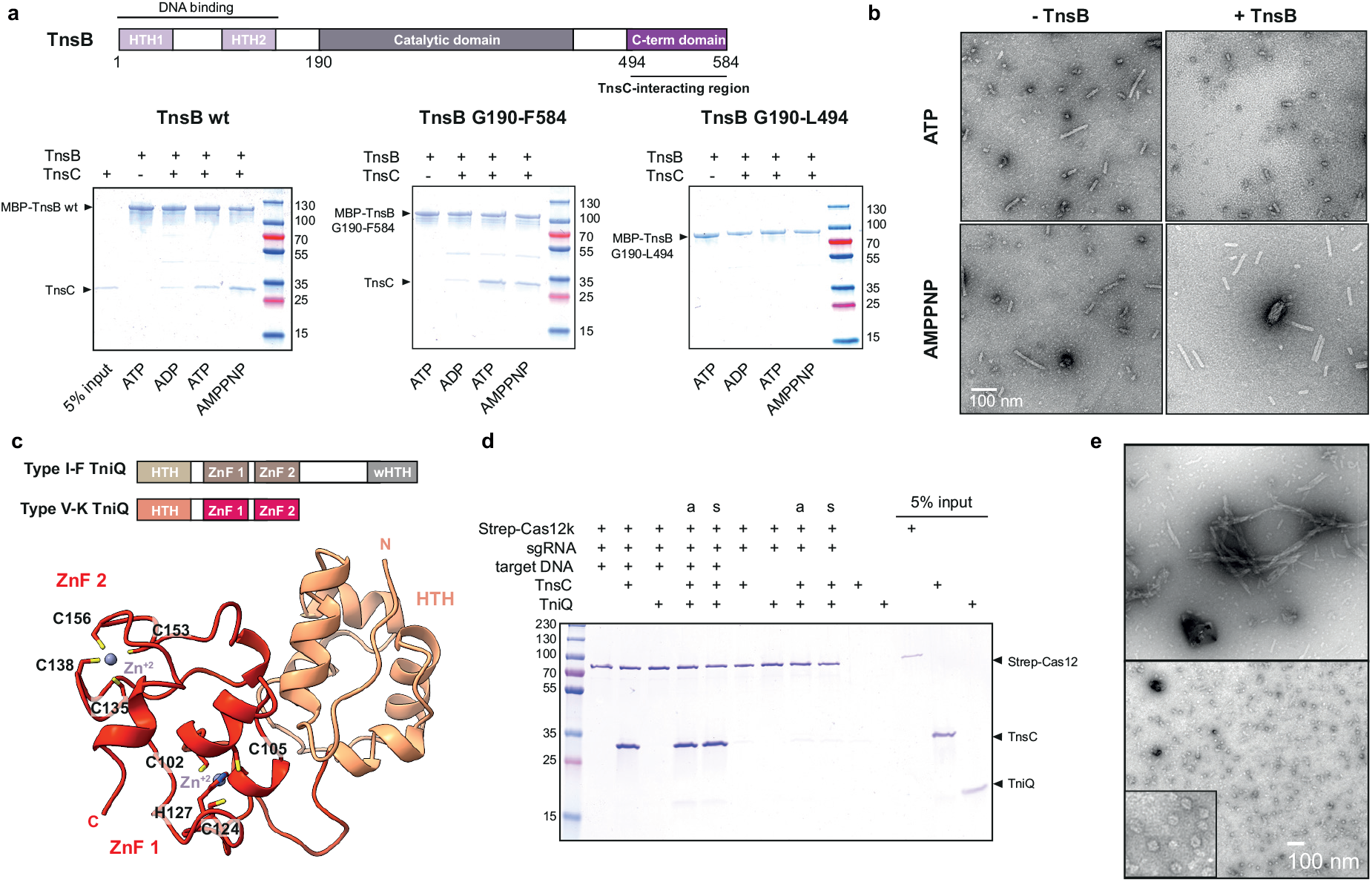
Functional interactions and roles of ShCAST components. **a**, Domain organization of ShTnsB (top panel). ZnF, zinc finger; HTH, helix turn helix motif. Co-precipitation of ShTnsC by ShTnsB constructs fused to maltose binding protein (MBP) in the presence of different nucleotides (bottom panel). **b**, Representative negative stain EM micrographs of ShTnsC-dsDNA complexes in the presence of ShTnsB and different nucleotides. Scale bars, 100 nm. Magnification, 98,000 x. **c**, Domain architecture of TniQ proteins (top panel). HTH, helix turn helix motif. wHTH, winged helix turn helix motif. ZnF, zinc-finger motif. Crystal structure of ShTniQ at 1.3 Å resolution (bottom panel). Side chains of zinc-coordinating residues are shown as sticks. Zinc ions are depicted as purple spheres. **d**, Co-precipitation of ShTnsC and ShTniQ by immobilized StrepII-fused Cas12k-sgRNA complex. a, TnsC and TniQ components were added together; s, sequential addition. **e**, Representative negative stain EM micrographs of ShTnsC in the presence of dsDNA, ATP and TniQ. Scale bars, 100 nm. Magnification of 68,000 x for main micrographs, 120,000 x for insert.

### The Zn finger protein TniQ restricts TnsC polymerization

TniQ is a putative DNA binding protein required for the DNA insertion activities of type V CRISPR-associated transposons *in vivo^7^*. In type I-F CRISPR-transposon systems, TniQ forms a head-to-tail dimer and interacts directly with the crRNA-guided Cascade effector complex^20–23^, but whether TniQ plays an analogous role in type V systems is not known. To obtain structural insights into TniQ function in type V systems, we initially determined the crystal structure of ShTniQ at a resolution of 1.3 Å (**Supplementary Table 3, Fig. 5c**). The structure reveals a conserved domain architecture comprising an N-terminal helix-turn-helix (HTH) motif followed by two zinc-finger domains (**Fig. 5c**), previously observed in the structure of TniQ from the *Vibrio cholerae* type I-F CRISPR-associated transposon^20–23^ (**Extended Data Fig. 10a**). However, unlike the dimeric *V. cholerae* TniQ, ShTniQ lacks the C-terminal winged HTH (wHTH) domain involved in dimerization, which is consistent with the observation that ShTniQ is monomeric in solution (**Extended Data Fig. 10b**). We tested whether TniQ associates with Cas12k to mediate interactions between Cas12k and TnsC using pull-down assays. We assembled StrepII-tagged ShCas12k with a sgRNA and a target DNA and immobilized the resulting complex on Strep-tactin beads. TnsC was efficiently co-precipitated by the Cas12k-sgRNA-target DNA complex in the presence of ATP, but did not bind in the absence of target DNA (**Fig. 5d**), indicating that the interaction is a result of ATP-dependent ShTnsC oligomerization on the target DNA. The interaction of ShTnsC with Cas12k was neither enhanced nor inhibited by the addition of ShTniQ (**Fig. 5d**), implying that TniQ is not required for TnsC assembly on target DNA, which corroborates previous observations that the RNA-guided DNA transposition activity of ShCAST is not dependent on TniQ *in vitro^7^*. Furthermore, ShTniQ was not co-precipitated by the Cas12k complex in the absence of TnsC (**Fig. 5d**), indicating that unlike in type I CRISPR-transposon systems, TniQ does not directly interact with the RNA-guided CRISPR effector module in type V-K systems. Finally, to probe whether TniQ modulates the DNA-induced oligomerization of TnsC, we examined ShTnsC filament formation in the presence of ShTniQ. Unexpectedly, addition of ShTniQ led to reduced ShTnsC polymerization (**Fig. 5e**), yielding substantially shorter filaments or ring-shaped assemblies, indicating that ShTniQ restricts ShTnsC polymerization. Taken together, these findings thus suggest that TniQ plays a distinct functional role in type V CRISPR-associated transposon systems.

## DISCUSSION

In this study, we provide structural and mechanistic insights into target DNA selection by Cas12k and subsequent remodeling by the transposon AAA+ ATPase TnsC in type V-K CRISPR-associated transposon systems. The ternary Cas12k complex structure reveals an extensive tracrRNA scaffold that appears to functionally compensate for the compact size of Cas12k, reminiscent of other small type V CRISPR-associated effectors, notably Cas12e (CasX)^30^. The structure reveals the atomic details of PAM recognition and R-loop formation, shedding light on the mechanism of RNA-guided DNA recognition. Notably, the Cas12k complex does not form a complete R-loop, drawing parallels with the Cascade-TniQ complex from I-F CRISPR-transposon systems and implying additional conformational activation steps by downstream transposon factors that may serve an activation checkpoint^20^. Furthermore, the R-loop structure is compatible with the reported minimal requirements for guide RNA-target complementarity in ShCAST^6^. The structure of TnsC filaments shows that ATP-dependent TnsC polymerization occurs on remodeled, underwound DNA, hinting that DNA unwinding and R-loop formation by Cas12k may help nucleate TnsC filament formation, although R-loop formation alone is not sufficient for ShCAST activity^7^. Based on our findings, we propose a mechanistic model of Cas12k-dependent DNA transposition in type V-K CRISPR-transposon systems, in which target site recognition by Cas12k initiates local TnsC polymerization to generate a recruitment platform for the transposase TnsB (**Fig. 6**). TnsB in turn stimulates the ATPase activity of TnsC to trigger filament disassembly, thereby exposing the transposon insertion site. Unlike in type I CRISPR-transposon systems, TniQ plays a fundamentally different role in type V-K systems by restricting TnsC polymerization to the vicinity of Cas12k, which is consistent with the lack of a direct Cas12k-TniQ interaction and with TniQ being dispensable for ShCAST-mediated DNA transposition *in vitro^7^*. Indeed, a complementary structural study of TnsC reveals that TniQ inhibits TnsC polymerization by capping TnsC filaments^31^. Altogether, this work discloses mechanistic insights into target site selection and remodelling by type V CRISPR-associated transposons and provides a structural framework for their development as an RNA-guided insertion technology for genome engineering.

**Figure 6.**
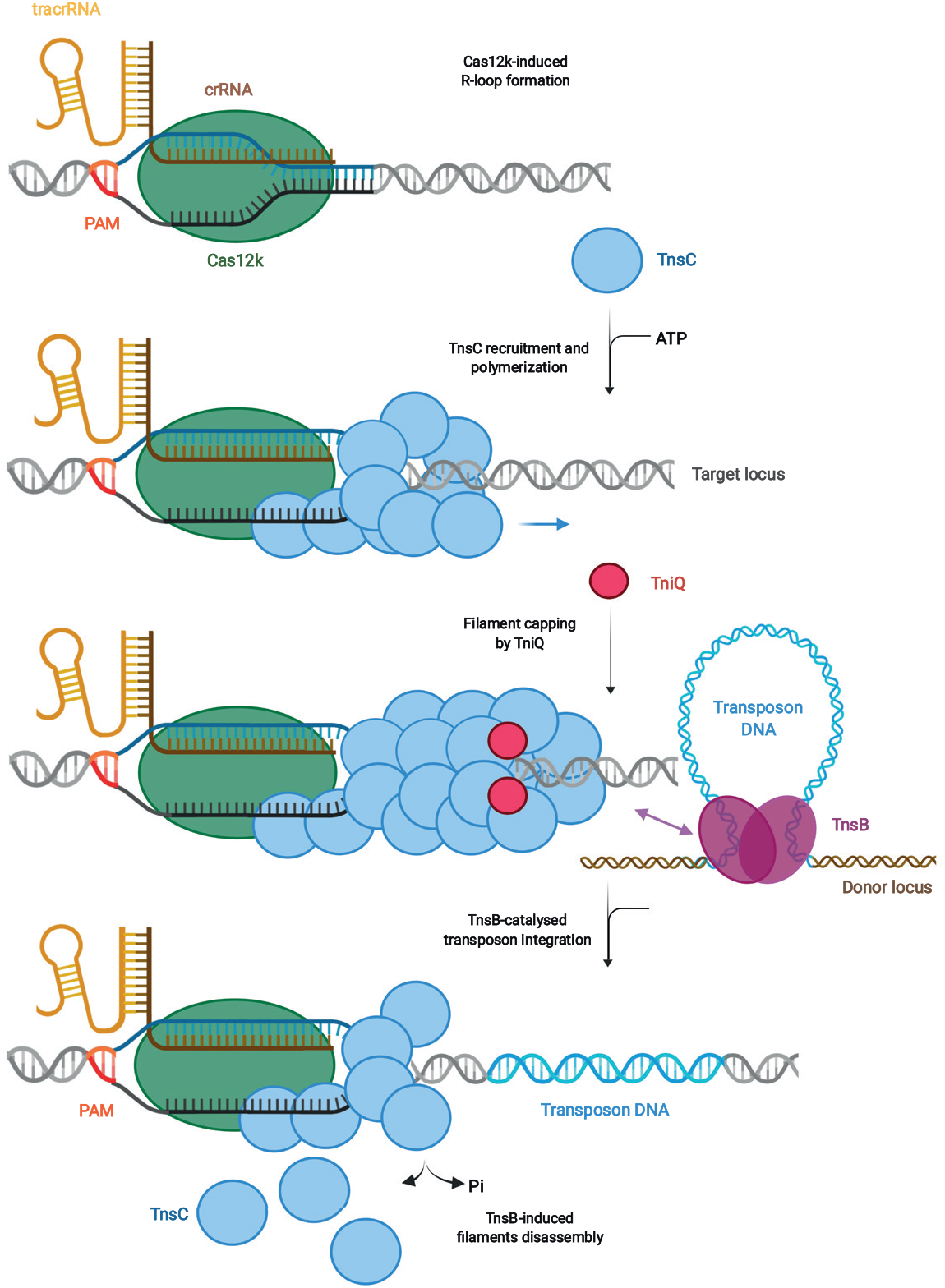
Mechanistic model of Cas12k-directed transposition. Schematic depicting transposition of type V-K CRISPR-associated transposons. Cas12k in association with a crRNA-tracrRNA dual guide RNA recognizes target DNA sequences, forming a partial R-loop structure. Full R-loop is formed upon recruitment of TnsC by interactions with DNA-bound Cas12k, which nucleates ATP-dependent formation of a helical filament around structurally remodeled DNA. Filament growth is restricted by TniQ capping the Cas12k-distal end. The TnsC filament serves as a recruitment platform for TnsB, which interacts directly with TnsC and stimulates its ATPase activity. This leads to filament disassembly and exposes the transposon insertion site in the DNA located at a fixed distance from Cas12k.

## METHODS

### Plasmid DNA constructs

The DNA sequences of *Scytonema hofmanni* Cas12k (WP_029636312.1), TnsC (WP_029636336.1), TniQ (WP_029636334.1) and TnsB (WP_084763316.1) proteins were codon optimized for heterologous expression in *Escherichia coli* (*E. coli*) and synthesized by GeneArt (Thermo Fisher Scientific). The ShCas12k gene was inserted into the 1B (Addgene 29653) and 1R (Addgene 29664) plasmids using ligation-independent cloning (LIC), resulting in constructs carrying an N-terminal hexahistidine tag followed by a tobacco etch virus (TEV) protease cleavage site and a N-terminal hexahistidine-StrepII tag followed by a TEV cleavage site, respectively. The ShTnsB gene and truncated derivatives were inserted by LIC into 1B (Addgene 29653) and 1C (Addgene 29659) plasmids, generating constructs carrying an N-terminal hexahistidine tag and a hexahistidine-maltose binding protein (MBP) tag followed by a TEV cleavage site. The ShTnsC gene was inserted into the 1S (Addgene 29659) plasmid to produce a construct carrying a N-terminal hexahistidine and hexahistidine-SUMO tag followed by a TEV cleavage site and the ShTniQ was cloned into a 1C vector. Point mutations were introduced by Gibson assembly using gBlock gene fragments synthetized by IDT or annealed oligonucleotides provided by Sigma as inserts. The pDonor (Addgene 127921), pHelper (Addgene 127924), and pTarget (Addgene 127926) plasmids^7^ used in droplet digital PCR experiments were sourced from Addgene. The PSP1-targeting spacer was cloned into pHelper by Gibson assembly, yielding pHelper-PSP1. Mutations in the sgRNA or in the sequence of the ShCas12k and ShTnsC genes were introduced into the pHelper plasmid by Gibson assembly. Plasmids were cloned and propagated in Mach I cells (Thermo Fisher Scientific) with the exception of pHelper, which was grown in One Shot PIR1 cells (Thermo Fisher Scientific). Plasmids were purified using the GeneJET plasmid miniprep kit (Thermo Fisher Scientific) and verified by Sanger sequencing. All sequence of primers, gBlocks and oligonucleotides used for cloning are provided in Supplementary Table 4.

### Protein expression and purification

For expression of ShCas12k constructs, hexahistidine-Strep II-tagged and hexahistidine-tagged ShCas12k proteins were expressed in *E. coli* BL21 Star (DE3) cells. Cell cultures were grown at 37 °C shaking at 100 rpm until reaching an OD_600_ of 0.6 and protein expression was induced with 0.4 mM IPTG (isopropyl-β-d-thiogalactopyranoside) and continued for 16 h at 18 °C. Harvested cells were resuspended in 20 mM Tris-HCl pH 8.0, 500 mM NaCl, 5 mM Imidazole, 1 μg/mL Pepstatin, 200 μg/mL AEBSF and lysed in a Maximator Cell homogenizer at 2000 bar and 4 °C. The lysate was cleared by centrifugation at 40,000 *g* for 30 min at 4 °C and applied to two 5mL Ni-NTA cartridges connected in tandem. The column was washed with 50 mL of 20 mM Tris-HCl pH 8.0, 500 mM NaCl, 5 mM Imidazole before elution with 25 mL of 20 mM Tris-HCl pH 8.0, 500 mM NaCl, 250 mM Imidazole. Elution fractions were pooled and dialyzed overnight against 20 mM HEPES-KOH pH 7.5, 250 mM KCl, 1 mM EDTA, 1 mM DTT in the presence of tobacco etch virus (TEV) protease. Dialyzed proteins were loaded onto a 5 mL HiTrap Heparin HP column (GE Healthcare) and eluted with a linear gradient of 20 mM HEPES-KOH pH 7.5, 1 M KCl. Elution fractions were pooled, concentrated using 30,000 molecular weight cut-off centrifugal filters (Merck Millipore) and further purified by size exclusion chromatography using a Superdex 200 (16/600) column (GE Healthcare) in 20 mM HEPES-KOH, 250 mM KCl, 1 mM DTT yielding pure, monodisperse proteins. Purified proteins were concentrated to 10-15 mg mL^−1^, flash frozen in liquid nitrogen and stored at −80 °C until further usage.

Expression of wild-type ShTnsC, ShTnsC mutants, ShTnsB, ShTnsB G190-F584, ShTnsB G190-L494 and ShTniQ was performed in *E. coli* BL21 Rosetta2 (DE3) cells. Cells were grown in LB medium until reaching an OD_600_ of 0.6 and expression was induced by addition of 0.4 mM IPTG. Proteins were expressed at 18 °C for 16 h. For ShTnsC, the cells were harvested and resuspended in lysis buffer containing 20 mM Tris-HCl pH 7.5, 500 mM NaCl, 5% glycerol and 10 mM imidazole supplemented with EDTA-free protease inhibitor (Roche). The cell suspension was lysed by ultrasonication and the lysate was cleared by centrifugation at 40,000 x *g* for 40 min. Cleared lysate was applied to a 5 mL Ni-NTA cartridge (Qiagen). The column was washed in two steps with lysis buffer supplemented with 25 and 100 mM imidazole, and bound proteins were eluted with 25 mL of same buffer supplemented with 500 mM imidazole pH 7.5. Eluted proteins were dialysed overnight against 20 mM Tris-HCl pH 7.5, 250 mM NaCl, 5% glycerol, 1 mM DTT in the presence of TEV protease. The proteins was further purified using a 5 mL HiTrap HP Heparin column (GE Healthcare) and eluted with a buffer containing 20 mM Tris-HCl pH 7.5, 700 mM NaCl, 5% glycerol and 1 mM DTT. The eluted fraction was concentrated and further purified by size exclusion chromatography using an S200 increase (10/300 GL) column (GE Healthcare) in 20 mM Tris-HCl pH 7.5, 500 mM NaCl, 1 mM DTT, yielding pure, monodisperse proteins. Purified ShTnsC was concentrated to 1-2 mg mL^−1^ using 30,000 molecular weight cut-off centrifugal filters (Merck Millipore) and flash-frozen in liquid nitrogen. All mutant ShTnsC proteins were purified using the same protocol as for the wild-type protein.

For ShTnsB, cells were harvested and resuspended in lysis buffer containing 20 mM Tris-HCl pH 8.0, 500 mM NaCl, 5% glycerol and 5 mM Imidazole supplemented with EDTA-free protease inhibitor (Roche) and lysed by ultrasonication. Lysate was clarified by centrifugation at 40,000 x *g* for 40 min. The lysate was applied to a 5 mL Ni-NTA cartridge (Qiagen) and the column was washed with lysis buffer supplemented with 25 mM imidazole. The protein was eluted with 20 mM Tris-HCl pH 7.5, 500 mM NaCl, 5% glycerol and 200 mM Imidazole. Eluted proteins were dialysed overnight against 20 mM Tris-HCl pH 8.0, 200 mM NaCl, 1 mM DTT in the presence of TEV protease and subsequently purified using a 5 mL HiTrap HP Heparin column (GE Healthcare), eluting with a buffer containing 20 mM Tris-HCl pH 8.0, 400 mM NaCl, and 1 mM DTT. The eluted fraction was concentrated and further purified by size exclusion chromatography using a Superdex (16/600) column (GE Healthcare) in 20 mM Tris-HCl pH 8.0, 400 mM NaCl, and 1 mM DTT. Purified ShTnsB was concentrated to 7 mg mL^−1^, and flash-frozen in liquid nitrogen. For the ShTnsB G190-F584 and ShTnsB G190-L494 constructs, cells were harvested and resuspended in 20 mM Tris-HCl pH 7.5, 500 mM NaCl, 5% glycerol and 5 mM imidazole supplemented with EDTA-free protease inhibitor (Roche) and lysed by ultrasonication. Cleared lysate was applied to a 5 mL Ni-NTA cartridge (Qiagen) and the column was washed in two steps with lysis buffer supplemented with 25 and 50 mM imidazole. The proteins were eluted with 20 mM Tris-HCl pH 7.5, 500 mM NaCl, 5% glycerol and 125 mM imidazole and dialysed overnight against 20 mM Tris-HCl pH 7.5, 500 mM NaCl, 5% glycerol and 4 mM beta-mercaptoethanol in the presence of TEV. Dialysed proteins were passed through a 5 mL Ni-NTA cartridge and washed with buffer containing 20 mM Tris-HCl pH 7.5, 500 mM NaCl, 5% glycerol and 25 mM imidazole, and further purified by size exclusion chromatography using an Superdex (16/600) column (GE Healthcare) in 20 mM Tris-HCl pH 7.5, 250 mM NaCl, 1 mM DTT. Purified ShTnsB proteins were concentrated to 13–24 mg mL^−1^ and flash-frozen in liquid nitrogen. For the purification of the MBP-tagged constructs used in pull-downs experiments (MBP-ShTnsB, MBP-ShTnsB G190-F584 and MBP-ShTnsB G190-L494), eluted fraction were set aside after the first step of Ni affinity chromatography and directly loaded on a Superdex 200 Increase (10/300 GL) column (GE Healthcare) and eluted in 20 mM Tris-HCl pH 7.5, 250 mM NaCl and 1 mM DTT. Appropriate fractions were concentrated to 3-0.7 mg mL^−1^ using 30,000 molecular weight cut-off centrifugal filters (Merck Millipore) and flash-frozen in liquid nitrogen.

For ShTniQ, cells were harvested and resuspended in 20 mM Tris-HCl pH 7.8, 500 mM NaCl, 5% glycerol and 5 mM imidazole supplemented with EDTA-free protease inhibitor (Roche) and lysed by ultrasonication. The cleared lysate was applied to a 5 mL Ni-NTA cartridge (Qiagen) and the column was washed in two steps with lysis buffer supplemented with 25 and 50 mM imidazole. The protein was eluted with lysis buffer supplemented with 300 mM imidazole. Eluted protein was dialysed overnight against 20 mM Tris-HCl pH 7.8, 500 mM NaCl, 1 mM DTT in the presence of TEV protease. Dialysed protein was passed through a 5 mL MBPTrap column (GE Healthcare). The flow-through fraction was concentrated and further purified by size exclusion chromatography using a Superdex 200 (16/600) column (GE Healthcare) in 20 mM Tris-HCl pH 7.8, 250 mM NaCl, 1 mM DTT. Size exclusion chromatography allowed to estimate the oligomeric state of the protein (monomer), as shown in Extended Data Figure S10b. Purified ShTniQ was concentrated to 10 mg mL^−1^ using 10,000 molecular weight cut-off centrifugal filters (Merck Millipore) and flash-frozen in liquid nitrogen.

### sgRNA preparation

DNA sequence encoding the T7 RNA polymerase promoter upstream of the ShCas12k-sgRNA was sourced as a gBlock (IDT), cloned into a pUC19 plasmid using restriction digest with BamHI & EcoRI, and confirmed by Sanger sequencing. The sequence encoding the T7 RNA polymerase promoter and sgRNA was amplified by PCR and purified by ethanol precipitation for use as template for *in vitro* transcription with T7 RNA polymerase as described previously^32^. The transcribed RNA was gel purified, precipitated with 70 % (v/v) ethanol, dried and dissolved in nuclease-free water.

### TnsC filament formation

For the reconstitution of ShTnsC filaments, wild-type ShTnsC protein was diluted to a final concentration of 0.5 μM in a buffer containing 20 mM HEPES pH 7.5, 200 mM KCl, 10 mM MgCl_2_, 1 mM DTT and mixed at a ratio of 1:5 with a double-stranded 92 bp duplex oligo (**Supplementary Table 4**) in a 24 μl reaction volume. Protein and DNA were incubated for 10 min at 25 °C. Upon addition of nucleotide (ATP, ATPγS or AMPPNP) at a final concentration of 1 mM, reactions were further incubated for 20 min at 37 °C. To analyse filament assembly in the presence of TnsB, TnsC filaments were formed using the same procedure described above, followed by addition of TnsB at a final concentration of 10 μM (1:2 ratio TnsC:TnsB) in a 60 μl final reaction volume. To probe filament formation in the presence of ShTniQ, ShTnsC was diluted to a final concentration of 3 μM in a buffer containing 20 mM HEPES pH 7.5, 200 mM KCl, 10 mM MgCl_2_, 1 mM DTT and 1 mM ATP and mixed at a ratio of 1:25 with a double-stranded 69 bp duplex oligo (**Supplementary Table 4**) in a 20 μl reaction volume. ShTniQ was then added to the reactions at a final concentration of 6 μM and incubated for 1 hour at 37 °C. Prepared samples were then analysed by negative stain electron microscopy.

### Negative stain electron microscopy

Samples (4 μL) were applied to glow-discharged continuous carbon film supported copper grids (CF300-CU-50 grids, Electron Microscopy Sciences). After 1 min incubation, the grids were washed once with 4 μl of 2 % (w/v) uranyl formate and stained in a 4 μL drop of 2 % uranyl formate during a total of 1 min prior blotting. Negatively-stained specimens were imaged using a FEI Tecnai G2 Spirit transmission electron microscope operated at an acceleration voltage of 120 kV. Data were acquired at a nominal magnification of 68,000, 98,000, 120,000 or 180,000 x, as indicated in figure legends. The length of the filaments was measured manually with the Gatan Digital micrograph software.

### Transposition assay and droplet digital PCR analysis

Transposition assay were conducted as in^6^, with minor modifications. All transposition experiments were performed in One Shot PIR1 *E. coli* cells (Thermo Fisher Scientific). The strain was first co-transformed with 20 ng each of pDonor and pTarget, and transformants were isolated by selective plating on double antibiotic LB-agar plates. Liquid cultures were then inoculated from single colonies, and the resulting strains were made chemically competent using standard methods, aliquoted and snap frozen. pHelper plasmids (20 ng) were then introduced in a new transformation reaction by heat shock, and after recovering cells in fresh LB medium at 37 °C for 1 h, cells were plated on triple antibiotic LB-agar plates containing 100 μg mL^−1^ carbenicillin, 50 μg mL^−1^ kanamycin, and 33 μg mL^−1^ chloramphenicol. After overnight growth at 37 °C for 16 h, colonies were harvested from the plates, resuspended in 15 μL Lysis buffer (TE with 0.1 % Triton X 100) and heated for 5 min at 95 °C. 60 μL of water were added to the samples before centrifugation for 10 min at 16,000 x *g*. The supernatant was then transferred and the nucleic acid concentration adjusted to 0.3 ng/μL. 0.75 ng of template DNA were used for subsequent investigation by droplet digital PCR (ddPCR). A mixture of five primers (900 nM final concentration each), two probes (250 nM final each) (**Supplementary Table 4**) and template DNA were combined with ddPCR Supermix for Probes (No dUTP) (BioRad) in a final 20 μL reaction volume. Droplets were generated with 70 μL of Droplet Generation Oil for Probes (BioRad) using the QX200 Droplet Generator (BioRad). 40 μL of final sample were transferred to 96 well plates for amplification by PCR (1 cycle, 95 °C, 10 min; 40 cycles, 94 °C, 30 s, 58 °C, 1 min; 1 cycle, 98 °C, 10 min; 4 °C hold). Samples were read with a QX200 Droplet Reader, and data analysed using the QuantaSoft (v1.6.6.0320) to determine the concentration of inserts and template in each reaction (Abs counting mode). On-target transposition efficiencies were calculated as inserts/(inserts+target). All ddPCR measurements presented in the text and figures were determined from three independent biological replicates and measured in technical duplicates.

### Pull-down experiments

For pull-down experiments using ShCas12k as bait, sgRNA was first mixed with hexahistidine-StrepII-tagged ShCas12k (Strep-Cas12k) in assembly buffer (20 mM HEPES pH 7.5, 250 mM KCl, 10 mM MgCl_2_, 1 mM DTT), and incubated 20 min at 37 °C to allow complex formation. A dsDNA target (**Supplementary Table 4**) was then added to the reaction in a Strep-Cas12k:sgRNA:dsDNA molar ratio of 1:1.2:2 and incubated for 20 min at 37 °C. The final 20 μL reaction contained 2.5 μg (at a concentration of 1.6 μM) Strep-Cas12k, 1.9 μM sgRNA, and 3.2 μM DNA in assembly buffer. Samples were mixed with 12.5 μL Strep-Tactin beads (iba) equilibrated in pull-down wash buffer 1 (20 mM HEPES pH 7.5, 250 mM KCl, 10 mM MgCl_2_, 1 mM DTT, 0.05 % Tween20) and incubated 20 min at 4 °C on a rotating wheel. The beads were washed three times with pull-down wash buffer 1 to remove excess nucleic acids. The beads were resuspended in 250 μL of pull-down wash buffer 1 and ShTniQ and/or ShTnsC was added in 10-fold molar excess together with AMPPNP (1 mM final concentration). Samples were incubated 20 min at 37 °C, then washed twice with pull-down wash buffer 2 (20 mM HEPES pH 7.5, 250 mM KCl, 10 mM MgCl_2_, 1 mM DTT, 0.05 % Tween20, 1 mM AMPPNP) and eluted with pull-down elution buffer (20 mM HEPES pH 7.5, 250 mM KCl, 10 mM MgCl_2_, 1 mM DTT, 1 mM AMPPNP, 5 mM desthiobiotin). Eluted samples were analyzed by SDS-PAGE using Any kDa gradient polyacrylamide gels (Bio-Rad) and stained with Coomassie Brillian Blue.

For pull-downs using ShTnsB as bait, MBP-ShTnsB and the truncated derivatives ShTnsB G190-F584 and ShTnsB G190-L494 (62.5 μmol) were immobilized on 25 μl amylose agarose beads (New England Biolabs) in pull-down buffer (20 mM HEPES pH 7.5, 200 mM KCl, 10 mM MgCl2, 1 mM DTT) by incubation for 30 min at 4 °C. The beads were resuspended and washed three times with 200 μL of pull-down buffer. In parallel, to allow filament formation, TnsC was first diluted to a final concentration of 8 μM in pull-down buffer and mixed at a molar ratio of 1:5 with a double-stranded 92-bp DNA duplex (**Supplementary Table 4**) in a 24 μL reaction volume and incubated for 10 min at 25 °C. Upon addition of nucleotide (ADP, ATP, or AMPPNP) at a final concentration of 1 mM, the 25 μL reactions were further incubated for 20 min at 37 °C. The 25 μL mixture containing ShTnsC was then added to the amylose beads containing TnsB as bait, followed by incubation for 15 min at 37 °C. Beads were then washed three times with 200 μL of pull-down buffer. Proteins were eluted from the beads by adding SDS loading buffer, and analyzed on 4–15% gradient polyacrylamide gel (Bio-Rad).

### DNA binding assays

Increasing concentrations of wild-type or mutant ShTnsC (0 nM, 1.3 μM, 2.5 μM, 8 μM) were incubated with a fluorescently labeled double-stranded 27 bp duplex oligonucletide (20 nM final concentration) (**Supplementary Table 4**) for 30 min at 37 °C in a buffer containing 20 mM HEPES pH 7.5, 200 mM KCl, 10 mM MgCl2, 1 mM DTT and 1 mM AMPPNP, in a final volume of 20 μL. Reactions were resolved on a 4–20% native polyacrylamide gel (Bio-Rad) and imaged using a Typhoon scanner (GE Helthcare).

### Crystallization and structure determination of TniQ

Crystals of ShTniQ were grown using the hanging drop vapor diffusion method. ShTniQ at a concentration of 6.3 mg mL^−1^ was mixed in a 1:1 ratio (1 μL + 1 μL) with reservoir solution containing 100 mM HEPES pH 7.5, 17–25 % PEG6000 and 250 mM NaCl. Crystals were cryoprotected with reservoir solution supplemented with 25 % (v/v) ethylene glycol and flash-cooled in liquid nitrogen. Data were collected at a temperature of 100 K at the beamline PXIII of the Swiss Light Source (Paul Scherrer Institute, Villigen, Switzerland) and processed with autoPROC^33^. The crystals belonged to space group *P*2_1_2_1_2_1_ and contained four copies of ShTniQ per asymmetric unit. Experimental phases were obtained by the single-wavelength anomalous diffraction (SAD) method from data measured at a wavelength of 1.0000 Å, utilizing the anomalous signal of Zn^+2^ scatters present in the protein. Initial phases were calculated using Phenix^34^ (Autosol^35^) and density modification of experimental phases in AutoSHARP^36^ resulted in an interpretable map which was used for initial model building using ARP/wARP^37^. The atomic model was completed by iterative building in Coot^38^ and refined with Phenix.refine^39^ (**Supplementary Table 3**).

### ShCas12k-sgRNA-DNA ternary complex sample preparation and cryo-EM data collection

The target strand and non-target strand DNAs were sourced as synthetic, HPLC-purified oligonucleotides (IDT) and annealed. sgRNA was pre-incubated with ShCas12k for 20 min at 37 °C in assembly buffer (20 mM HEPES pH 7.5, 50 mM KCl, 10 MgCl_2_, 1 mM DTT). dsDNA was added and the sample was further incubated for 20 min at 37 °C. The final sample contained 47.8 μM ShCas12k, 57.4 μM sgRNA and 334.6 μM dsDNA (1:1.2:7 molar ratio) in 20 mM HEPES pH 7.5, 50 mM KCl, 10 mM MgCl_2_, 1 mM DTT. The sample was purified by size exclusion chromatography using a Superdex 200 (5/150) column. 2.5 μL of selected elution fractions were applied to a 300-mesh holey carbon grid (Au 1.2/1.3, Quantifoil Micro Tools), blotted 2 s at 75 % humidity, 4 °C, plunge frozen in liquid ethane (Vitrobot, FEI) and stored in liquid nitrogen. Cryo-EM data collection was performed on a FEI Titan Krios microscope operated at 300 kV equipped with a Gatan K3 direct electron detector using counting mode (EMBL Heidelberg, Germany). A total of 10,952 micrographs were recorded at a calibrated magnification of 130,000 x with a pixel size of 0.64 Å. Each movie comprises 40 subframes with a total dose of 51.81 e− Å^−2^. Data acquisition was performed automatically using SerialEM^40^ with four shots per hole at −0.9 μm to −1.9 μm defocus (0.1 μm steps).

### Image processing, model building for Cas12k-sgRNA-DNA complex

ShCas12k:sgRNA:dsDNA data was processed using cryoSPARC^41^. 10,952 movies were imported and motion-corrected (patch motion correction (multi)). Upon patch CTF estimation (multi), exposures were selected by estimated resolution (better than 3.5 Å) and defocus (<2 μm), yielding 10,866 movies. Particles were picked on denoised micrographs using Topaz^42^ with the pretrained model ResNet16 (64 units). Particle picks were inspected and selected particles (3.3 million) extracted in a box size of 300 × 300 pixel (fourier cropped to 100 × 100 pixel) and subjected to 2D classification. Particle-containing classes were selected (101 classes, 2.2 mio particles) and rebalanced to 6 super-classes each containing a maximum of 200,000 particles, yielding a final set of 1.2 million particles. The particle stack was used to calculate *ab initio* models. Two *ab initio* models matching the appearance of 2D classes were selected as volumes for heterogeneous refinement with the particle stack, resulting in eight 3D classes. Classes were inspected visually, respective particles re-extracted in unbinned form pixels and subjected to non-uniform refinement using the respective size-corrected volume from the heterogeneous refinement as input. The local resolution was calculated based on the resulting map using the local resolution functionality and plotted on the map using UCSF Chimera^43^. The structure model for the ShCas12k:sgRNA:dsDNA was built in Coot^38^. The sgRNA and target DNA were built *de novo* exclusively. The model was refined in Coot using restraints for the nucleic acids calculated with LIBG^44^(base pair, stacking plane and sugar pucker restraints) and finally refined using Phenix^34,39^. Real space refinement was performed with the global minimization and atomic displacement parameter (ADP) refinement options selected. Secondary structure restraints, side chain rotamer restraints, and Ramachandran restraints were used. Key refinement statistics are listed in **Supplementary Table 1**. The quality of the atomic models, including basic protein and DNA geometry, Ramachandran plots, and clash analysis, was assessed and validated with MolProbity^45,46^ in Phenix.

### ShTnsC-dsDNA complex cryo-EM sample preparation and data collection

Purified ShTnsC was diluted to a final concentration of 6.4 μM in a buffer containing 20 mM HEPES pH 7.5, 200 mM KCl, 10 mM MgCl_2_, 1 mM DTT and mixed at a ratio of 1:5 with a 92 nt dsDNA oligonucleotide containing two mismatched regions (**Supplementary Table 4**) in a 21.4 μL reaction volume and incubated for 10 min at 25 °C. Upon addition of AMPPNP at a final concentration of 1 mM, the sample was further incubated for 20 min at 37 °C. Sample vitrification was performed with a Leica EM GP2 plunge freezer at 10 °C and at 70 % humidity. 3.5 μL sample was applied onto a freshly glow-discharged 200-mesh copper 2 nm C R1.2/1.3 grids (Quantifoil Micro Tools). The grids were blotted for 1 s and plunge-vitrified in liquid ethane. Cryo-EM data collection was performed on a FEI Titan Krios G3i microscope (University of Zurich, Switzerland) operated at 300 kV equipped with a Gatan K3 direct electron detector in super-resolution counting mode. A total of 5,055 micrographs were recorded at a calibrated magnification of 130,000 × resulting in super-resolution pixel size of 0.325 Å. Each movie comprised 36 subframes with a total dose of 66.39 e− Å^−2^. Data acquisition was performed with EPU Automated Data Acquisition Software for Single Particle Analysis (ThermoFisher) with three shots per hole at −1.0 μm to −2.4 μm defocus (0.2 μm steps).

### Image processing, model building and model validation for ShTnsC-dsDNA complex

All movies were motion-corrected and dose-weighted with MOTIONCOR2^47^. Aligned, non-dose-weighted micrographs were then used to estimate the contrast transfer function (CTF) with GCTF^48^. All subsequent image processing steps were performed using helical reconstruction in RELION 3.1.2^49,50^. Approximately 1,100 segments were manually picked in RELION. One round of reference-free 2D classification was performed to produce templates for reference-dependent auto-picking. A limited resolution E-step (low-pass filter) of 10 Å was applied to prevent overfitting. Using the resulting 2D classes as templates, overlapping helical filament segments were automatically picked with an inter-box distance of 10 Å. A total of 1,733,884 segments were extracted with a box size of 352 pixels from the full dataset and subjected to three cycles of reference-free 2D classification to remove low-quality filaments, yielding 743,027 particles. Three iterative cycles of 3D auto-refinement in RELION using a cylinder with a 12 nm diameter as the initial model were conducted to produce initial reconstructions. A spherical mask with a diameter equal to 90 % of the box size was applied. Selected 2D classes were then used with the best *ab initio* model to perform 3D auto-refinement with a local search of helical symmetry (twist 58°–62° and rise 6–8 Å). The helical parameters were estimated based on a pitch of 41 Å calculated from the power spectra and assuming a number of 6 units/turns. The central Z-height parameter in RELION was set to 30 %. The output volume (3.94 Å resolution) was used as the reference model for 3D classification. The particles were divided into four classes in 3D classification. A 3D class with the most clearly discernible secondary structure features (59.44° twist and 6.65 Å rise and 77,544 particles) was selected and 3D auto-refined. A single round of Bayesian polishing, further helical refinement and post-processing with a soft mask applied around the filament produced a volume with final reported resolution of 3.57 Å, final twist 59.72° and rise 6.78 Å. The 3D reconstruction was sharpened with a B-factor of −59.51 Å^−2^. The overall map resolution reported in **Supplementary Table 2** was derived from Fourier shell correlation (FSC) calculations between reconstructions from two independently refined half-maps, and reported resolutions are based on the gold-standard criterion. Local resolution was estimated with RELION. For model building, a TnsC homology model based on the crystal structure of the DNA replication initiator factor ORC2 (PDB:1W5T)^51^ was generated using the Phyre2 server^52^ and the two main domains (α/β domain and C-terminal domain) were manually docked separately as rigid bodies in the cryo-EM density map with UCSF Chimera^43^, followed by real space fitting with the Fit in Map function. The atomic model of a ShTnsC protomer was manually rebuilt in Coot^38^. The coordinated AMPPNP molecules and magnesium ions were manually docked in Coot. The DNA duplex was manually built in Coot and then subjected to refinement in Coot using base pair, stacking plane and sugar pucker restraints generated in LIBG^44^. The sequence of the dsDNA was randomly selected and is purely representative as any DNA sequence information was lost during helical symmetry averaging. The adjacent subunits in the filament were then generated by applying the helical symmetry from RELION to the rebuilt atomic model. Real-space refinement of the resulting model was performed in Phenix^34,39^ with the global minimization and atomic displacement parameter (ADP) refinement options selected. The following restraints were used in real-space refinement: secondary structure restraints, non-crystallographic symmetry (NCS) restraints between the protein subunits, side chain rotamer restraints, and Ramachandran restraints. Key refinement statistics are listed in **Supplementary Table 2**. The quality of the atomic models, including basic protein and DNA geometry, Ramachandran plots, and clash analysis, was assessed and validated with MolProbity^45,46^ as implemented in Phenix.

## Data Availability

Atomic coordinates of the reported cryo-EM and X-ray crystallographic structures have been deposited, together with maps or structure factors with the Protein Data Bank under accession numbers 7OXD (ShTniQ).

## Acknowledgements

We are grateful to Marta Sawicka and Simona Sorrentino (University of Zurich Center for Microscopy and Image Analysis) for assistance with cryo-EM sample preparation and data collection. We acknowledge the Cryo-Electron Microscopy Service Platform at EMBL Heidelberg for instrument access and are grateful to Felix Weis for assistance with data collection. We a grateful to Martin Pacesa for help with cryo-EM data processing and Franziska Boneberg for technical assistance. We thank Beat Blattmann at the Protein Crystallization Center (University of Zurich) for assistance with crystallization screening; Meitian Wang, Vincent Olieric, and Takashi Tomizaki at the Swiss Light Source (Paul Scherrer Institute, Villigen, Switzerland) for assistance with X-ray diffraction measurements. We thank Susanne Kreutzer and the ETH Genome Engineering and Measurement Lab for assistance with ddPCR assays. This work was supported by Swiss National Science Foundation Project Grant 31003A_182567 and European Research Council (ERC) Consolidator Grant no. ERC-CoG-820152. I.Q. was supported by FEBS and EMBO (ALTF 296-2020) long-term postdoctoral fellowships. M.J. is an International Research Scholar of the Howard Hughes Medical Institute, and Vallee Scholar of the Bert L & N Kuggie Vallee Foundation.

## Author Contributions

I.Q., M.S. and M.J. conceived the study and designed experiments. I.Q. determined the structure of ShTnsC. M.S. conducted cryo-EM analysis of Cas12k. M.S. and I.Q. carried out biochemical and ddPCR functional experiments and negative-stain EM analysis. S.O. expressed and purified ShTnsC mutant proteins. C.C. assisted with sample preparation for biochemical and ddPCR assays. I.Q., M.S. and M.J. analyzed the data and wrote the manuscript.

## REFERENCES

1 Sorek, R., Lawrence, C. M. & Wiedenheft, B. CRISPR-mediated adaptive immune systems in bacteria and archaea. Annu Rev Biochem 82, 237–266, doi:10.1146/annurev-biochem-072911-172315 (2013).

2 Faure, G. et al. CRISPR-Cas in mobile genetic elements: counter-defence and beyond. Nat Rev Microbiol 17, 513–525, doi:10.1038/s41579-019-0204-7 (2019).

3 Klompe, S. E., Vo, P. L. H., Halpin-Healy, T. S. & Sternberg, S. H. Transposon-encoded CRISPR-Cas systems direct RNA-guided DNA integration. Nature, doi:10.1038/s41586-019-1323-z (2019).

4 Petassi, M. T., Hsieh, S. C. & Peters, J. E. Guide RNA Categorization Enables Target Site Choice in Tn7-CRISPR-Cas Transposons. Cell 183, 1757–1771 e1718, doi:10.1016/j.cell.2020.11.005 (2020).

5 Peters, J. E., Makarova, K. S., Shmakov, S. & Koonin, E. V. Recruitment of CRISPR-Cas systems by Tn7-like transposons. Proc Natl Acad Sci U S A 114, E7358–E7366, doi:10.1073/pnas.1709035114 (2017).

6 Saito, M. et al. Dual modes of CRISPR-associated transposon homing. Cell, doi:10.1016/j.cell.2021.03.006 (2021).

7 Strecker, J. et al. RNA-guided DNA insertion with CRISPR-associated transposases. Science, doi:10.1126/science.aax9181 (2019).

8 Aziz, R. K., Breitbart, M. & Edwards, R. A. Transposases are the most abundant, most ubiquitous genes in nature. Nucleic Acids Res 38, 4207–4217, doi:10.1093/nar/gkq140 (2010).

9 Chuong, E. B., Elde, N. C. & Feschotte, C. Regulatory activities of transposable elements: from conflicts to benefits. Nat Rev Genet 18, 71–86, doi:10.1038/nrg.2016.139 (2017).

10 Koonin, E. V., Makarova, K. S. & Wolf, Y. I. Evolutionary Genomics of Defense Systems in Archaea and Bacteria. Annu Rev Microbiol 71, 233–261, doi:10.1146/annurev-micro-090816-093830 (2017).

11 Peters, J. E. & Craig, N. L. Tn7: smarter than we thought. Nat Rev Mol Cell Biol 2, 806–814, doi:10.1038/35099006 (2001).

12 May, E. W. & Craig, N. L. Switching from cut-and-paste to replicative Tn7 transposition. Science 272, 401–404, doi:10.1126/science.272.5260.401 (1996).

13 Sarnovsky, R. J., May, E. W. & Craig, N. L. The Tn7 transposase is a heteromeric complex in which DNA breakage and joining activities are distributed between different gene products. EMBO J 15, 6348–6361 (1996).

14 Choi, K. Y., Spencer, J. M. & Craig, N. L. The Tn7 transposition regulator TnsC interacts with the transposase subunit TnsB and target selector TnsD. Proc Natl Acad Sci U S A 111, E2858–2865, doi:10.1073/pnas.1409869111 (2014).

15 Ronning, D. R. et al. The carboxy-terminal portion of TnsC activates the Tn7 transposase through a specific interaction with TnsA. EMBO J 23, 2972–2981, doi:10.1038/sj.emboj.7600311 (2004).

16 Stellwagen, A. E. & Craig, N. L. Gain-of-function mutations in TnsC, an ATP-dependent transposition protein that activates the bacterial transposon Tn7. Genetics 145, 573–585 (1997).

17 Mitra, R., McKenzie, G. J., Yi, L., Lee, C. A. & Craig, N. L. Characterization of the TnsD-attTn7 complex that promotes site-specific insertion of Tn7. Mob DNA 1, 18, doi:10.1186/1759-8753-1-18 (2010).

18 Parks, A. R. et al. Transposition into replicating DNA occurs through interaction with the processivity factor. Cell 138, 685–695, doi:10.1016/j.cell.2009.06.011 (2009).

19 Wolkow, C. A., DeBoy, R. T. & Craig, N. L. Conjugating plasmids are preferred targets for Tn7. Genes Dev 10, 2145–2157, doi:10.1101/gad.10.17.2145 (1996).

20 Halpin-Healy, T. S., Klompe, S. E., Sternberg, S. H. & Fernandez, I. S. Structural basis of DNA targeting by a transposon-encoded CRISPR-Cas system. Nature 577, 271–274, doi:10.1038/s41586-019-1849-0 (2020).

21 Jia, N., Xie, W., de la Cruz, M. J., Eng, E. T. & Patel, D. J. Structure-function insights into the initial step of DNA integration by a CRISPR-Cas-Transposon complex. Cell Res 30, 182–184, doi:10.1038/s41422-019-0272-2 (2020).

22 Li, Z., Zhang, H., Xiao, R. J. & Chang, L. F. Cryo-EM structure of a type I-F CRISPR RNA guided surveillance complex bound to transposition protein TniQ. Cell Res 30, 179–181, doi:10.1038/s41422-019-0268-y (2020).

23 Wang, B. B., Xu, W. H. & Yang, H. Structural basis of a Tn7-like transposase recruitment and DNA loading to CRISPR-Cas surveillance complex. Cell Res 30, 185–187, doi:10.1038/s41422-020-0274-0 (2020).

24 Li, Z. C., N.L. and Peters, J.E. in In: A.P. Roberts & P. Mullany (Eds.) Bacterial integrative mobile genetic elements. Austin, Texas: Landes Bioscience. 1–32 (2013).

25 Arias-Palomo, E. & Berger, J. M. An Atypical AAA+ ATPase Assembly Controls Efficient Transposition through DNA Remodeling and Transposase Recruitment. Cell 162, 860–871, doi:10.1016/j.cell.2015.07.037 (2015).

26 Mizuno, N. et al. MuB is an AAA+ ATPase that forms helical filaments to control target selection for DNA transposition. Proc Natl Acad Sci U S A 110, E2441–2450, doi:10.1073/pnas.1309499110 (2013).

27 Stellwagen, A. E. & Craig, N. L. Avoiding self: two Tn7-encoded proteins mediate target immunity in Tn7 transposition. EMBO J 16, 6823–6834, doi:10.1093/emboj/16.22.6823 (1997).

28 Greene, E. C. & Mizuuchi, K. Target immunity during Mu DNA transposition. Transpososome assembly and DNA looping enhance MuA-mediated disassembly of the MuB target complex. Mol Cell 10, 1367–1378, doi:10.1016/s1097-2765(02)00733-5 (2002).

29 Greene, E. C. & Mizuuchi, K. Dynamics of a protein polymer: the assembly and disassembly pathways of the MuB transposition target complex. EMBO J 21, 1477–1486, doi:10.1093/emboj/21.6.1477 (2002).

30 Liu, J. J. et al. CasX enzymes comprise a distinct family of RNA-guided genome editors. Nature 566, 218–223, doi:10.1038/s41586-019-0908-x (2019).

31 Park J., T. A., Mehrotra E., Petassi M.T., Hsieh S., Ke A., Peters J.E., Kellogg E.H., doi: bioRxiv doi: 10.1101/2019.12.11.123456 (2021).

32 Anders, C., Niewoehner, O. & Jinek, M. In Vitro Reconstitution and Crystallization of Cas9 Endonuclease Bound to a Guide RNA and a DNA Target. Methods Enzymol 558, 515–537, doi:10.1016/bs.mie.2015.02.008 (2015).

33 Vonrhein, C. et al. Data processing and analysis with the autoPROC toolbox. Acta Crystallogr D Biol Crystallogr 67, 293–302, doi:10.1107/S0907444911007773 (2011).

34 Liebschner, D. et al. Macromolecular structure determination using X-rays, neutrons and electrons: recent developments in Phenix. Acta Crystallogr D Struct Biol 75, 861–877, doi:10.1107/S2059798319011471 (2019).

35 Terwilliger, T. C. et al. Decision-making in structure solution using Bayesian estimates of map quality: the PHENIX AutoSol wizard. Acta Crystallogr D Biol Crystallogr 65, 582–601, doi:10.1107/S0907444909012098 (2009).

36 Vonrhein, C., Blanc, E., Roversi, P. & Bricogne, G. Automated structure solution with autoSHARP. Methods Mol Biol 364, 215–230, doi:10.1385/1-59745-266-1:215 (2007).

37 Langer, G., Cohen, S. X., Lamzin, V. S. & Perrakis, A. Automated macromolecular model building for X-ray crystallography using ARP/wARP version 7. Nat Protoc 3, 1171–1179, doi:10.1038/nprot.2008.91 (2008).

38 Emsley, P., Lohkamp, B., Scott, W. G. & Cowtan, K. Features and development of Coot. Acta Crystallogr D Biol Crystallogr 66, 486–501, doi:10.1107/S0907444910007493 (2010).

39 Afonine, P. V. et al. Real-space refinement in PHENIX for cryo-EM and crystallography. Acta Crystallogr D Struct Biol 74, 531–544, doi:10.1107/S2059798318006551 (2018).

40 de la Cruz, M. J., Martynowycz, M. W., Hattne, J. & Gonen, T. MicroED data collection with SerialEM. Ultramicroscopy 201, 77–80, doi:10.1016/j.ultramic.2019.03.009 (2019).

41 Punjani, A., Rubinstein, J. L., Fleet, D. J. & Brubaker, M. A. cryoSPARC: algorithms for rapid unsupervised cryo-EM structure determination. Nat Methods 14, 290–296, doi:10.1038/nmeth.4169 (2017).

42 Bepler, T. et al. Positive-unlabeled convolutional neural networks for particle picking in cryo-electron micrographs. Res Comput Mol Biol 10812, 245–247 (2018).

43 Pettersen, E. F. et al. UCSF Chimera--a visualization system for exploratory research and analysis. J Comput Chem 25, 1605–1612, doi:10.1002/jcc.20084 (2004).

44 Brown, A. et al. Tools for macromolecular model building and refinement into electron cryo-microscopy reconstructions. Acta Crystallogr D Biol Crystallogr 71, 136–153, doi:10.1107/S1399004714021683 (2015).

45 Chen, V. B. et al. MolProbity: all-atom structure validation for macromolecular crystallography. Acta Crystallogr D Biol Crystallogr 66, 12–21, doi:10.1107/S0907444909042073 (2010).

46 Prisant, M. G., Williams, C. J., Chen, V. B., Richardson, J. S. & Richardson, D. C. New tools in MolProbity validation: CaBLAM for CryoEM backbone, UnDowser to rethink “waters,” and NGL Viewer to recapture online 3D graphics. Protein Sci 29, 315–329, doi:10.1002/pro.3786 (2020).

47 Zheng, S. Q. et al. MotionCor2: anisotropic correction of beam-induced motion for improved cryo-electron microscopy. Nat Methods 14, 331–332, doi:10.1038/nmeth.4193 (2017).

48 Zhang, K. Gctf: Real-time CTF determination and correction. J Struct Biol 193, 1–12, doi:10.1016/j.jsb.2015.11.003 (2016).

49 He, S. & Scheres, S. H. W. Helical reconstruction in RELION. J Struct Biol 198, 163–176, doi:10.1016/j.jsb.2017.02.003 (2017).

50 Scheres, S. H. A Bayesian view on cryo-EM structure determination. J Mol Biol 415, 406–418, doi:10.1016/j.jmb.2011.11.010 (2012).

51 Singleton, M. R. et al. Conformational changes induced by nucleotide binding in Cdc6/ORC from Aeropyrum pernix. J Mol Biol 343, 547–557, doi:10.1016/j.jmb.2004.08.044 (2004).

52 Kelley, L. A., Mezulis, S., Yates, C. M., Wass, M. N. & Sternberg, M. J. The Phyre2 web portal for protein modeling, prediction and analysis. Nat Protoc 10, 845–858, doi:10.1038/nprot.2015.053 (2015).

53 Swarts, D. C. & Jinek, M. Mechanistic Insights into the cis- and trans-Acting DNase Activities of Cas12a. Mol Cell 73, 589–600 e584, doi:10.1016/j.molcel.2018.11.021 (2019).

54 Krissinel, E. & Henrick, K. Secondary-structure matching (SSM), a new tool for fast protein structure alignment in three dimensions. Acta Crystallogr D Biol Crystallogr 60, 2256–2268, doi:10.1107/S0907444904026460 (2004).

